# Simpati: patient classifier identifies signature pathways based on similarity networks for the disease prediction

**DOI:** 10.1101/2021.09.23.461100

**Authors:** Luca Giudice

## Abstract

**BACKGROUND:** Pathway-based patient classification is a supervised learning task which implies a model learning pathways as features to predict the classes of patients. The counterpart of enrichment tools for the pathway analysis are fundamental methods for clinicians and biomedical scientists. They allow to find signature cellular functions which help to define and annotate a disease phenotype. They provide results which lead human experts to manually classify patients. It is a paradox that pathway-based classifiers which natively resolve this objective are not strongly developed. They could simulate the human way of thinking, decipher hidden multivariate relationships between the deregulated pathways and the disease phenotype, and provide more information than a probability value. Instead, there are currently only two classifiers of such kind, they require a nontrivial hyperparameter tuning, are difficult to interpret and lack in providing new insights. There is the need of new classifiers which can provide novel perspectives about pathways, be easy to apply with different biological omics and produce new data enabling a further analysis of the patients.

**RESULTS:** We propose Simpati, an innovative and interpretable patient classifier based on pathway-specific patient similarity networks. The first classifier to adopt ad-hoc novel algorithms for such graph type. It standardizes the biological high-throughput dataset of patient’s profiles with a propagation algorithm that considers the interconnected nature of the cell’s molecules for inferring a new activity score. This allows Simpati to classify with dense, sparse, and non-homogenous omic data. Simpati organizes patient’s molecules in pathways represented by patient similarity networks for being interpretable, handling missing data and preserving the patient privacy. A network represents patients as nodes and a novel similarity measure determines how much every pair act co-ordinately in a pathway. Simpati detects signature biological processes based on how much the topological properties of the related networks separate the patient classes. In this step, it includes a new cohesive subgroup detection algorithm to handle patients not showing the same pathway activity as the other class members. An unknown patient is then classified by a unique recommender system which considers how much is similar to known patients and distant from being an outlier. Simpati outperforms previously published classifiers on five cancer datasets described with two biological omics, classifies well with sparse data, identifies more relevant pathways associated to the patient’s disease than the competitors and has the lowest computational requirements.

**CONCLUSION:** Simpati can serve as generic-purpose pathway-based classifier of patient classes. It provides signature pathways to unveil the altered biological mechanisms of a disease phenotype and to classify patients according to the learnt pathway-specific similarities. The signature condition and patient prediction can be deciphered considering the patient similarity networks which must reveal the members of a patient class more cohesive and similar than the non-members. Simpati divides the pathways in up and downinvolved. Upinvolved when the signaling cascades generated by the altered molecules of the disease patients impact stronger the pathway than the ones of the control class. We provide an R implementation, a graphical user interface and a visualization function for the patient similarity networks. The software is available at: https://github.com/LucaGiudice/Simpati

## 1. INTRODUCTION

High-throughput biological data provide valuable information to clinicians for the prognosis and treatment response of patients. They offer quantitative and qualitative evidences to biomedical scientists for developing a study or confirming wet-lab results [1–3]. Pathway-based analysis is a technique to interrogate these data. It provides an intuitive and comprehensive understanding of the molecular mechanisms related to the patients [4,5]. The pathway space is more robust to noise than the single feature level, summarizes the information of multiple patient’s molecules into the pathway activity (inhibited or activated), reduces the model complexity and maintains predictive accuracy in the face of uncontrolled variation [6–9]. For these reasons, it is strongly used to annotate a class of disease patients, to classify new patients and to find overlaps with old and novel findings. As example, the Kidney renal clear cell carcinoma (KIRC) patients exhibit high heterogeneity, the efficacy of therapies targeting single biomarkers greatly varies among patients. The research is currently considering consistent altered pathways as markers to recognise the disease and as targets to tackle the tumor at the earliest stage. The latter is currently associated to the upregulation of the innate immune system and inhibition of the cellular catabolism, while the tumor late stage is characterized by a strong upregulation of the cell adhesion, growth factors and hypoxia [10,11].

These motivations boosted the development of enrichment tools for the pathway analysis but not of machine learning algorithms. The latter are neither considered in reviews [12–15] nor in bioinformatics best practises [16–18]. Fabris et al. [19] detailed the drawbacks of a supervised classification approach. It lacks a formal statistical basis, is computationally expensive, includes a not trivial hyper-parameter setting and does not handle well neither imbalances classes nor structured feature types as the biological pathways.

Few attempts have been made in this direction. In 2010, Pang et al. [20] proposed a bivariate node-splitting random forest integrating pathways for the survival analysis of cancer patients in microarray studies. In 2018, Hao et al. [21] proposed the first generic-purpose pathway-based deep neural network for the prediction of Glioblastoma patients. The method builds a network model by leveraging prior biological knowledge of pathway databases and predicts considering hierarchical nonlinear relationships between the biological processes and the patient classes in comparison. However, the method requires a non-trivial tuning of hyper-parameters which are difficult to interpret for bioinformaticians or computational biologists without a background in deep learning, demands high computational resources, does not provide a graphical representation to explain why specific pathways have been selected and includes in the results only the classification performances.

In the same year, Pai et al. officially introduces the emerging patient similarity network (PSN) paradigm [22] for the precision medicine. In a PSN, each node is an individual patient and an edge between two patients corresponds to pairwise similarity for a given datum (gender, height, gene expression …). The paradigm brings many advantages. Analysing the similarities to gather new information is conceptually intuitive. A PSN can lead to the identification of patient subgroups or the prediction of a patient’s class. Similarity networks can represent any datum, naturally handle missing and heterogenous data, have a history of successes in gene and protein function prediction [23–25] and can preserve the patient privacy by being shared in place of the sensitive raw information (topic which is growing in concern) [26–28].

Pai et al. with our contribution propose netDx [29,30] as patient classifier based on the PSN paradigm. Any available datum (e.g., age, gender, gene expression, …) is converted into a PSN. The method then relies on GeneMANIA [25], state-of-art gene function predictor, to select the best patient similarity networks and to use them in the classification. GeneMANIA scores the PSNs based on how well classify training patients, creates a consensus network from the best scored PSNs, integrates the testing patients and predicts their class based on their similarity with the known ones. netDx has not been designed to be a pathway-based classifier but supports the creation of PSNs from gene-based data divided into user-defined groups as cellular functions. In this last scenario, it is possible to impose to netDx to create a PSN for each pathway. Despite researchers’ efforts for standing up to the challenge, the method does not accept data in matrix format, requires to define multiple functions in order to set up the model, relies on an external software to handle similarity networks, does not provide a graphical representation of the PSNs used to predict, does not give access to the data processed during the workflow, depends by the quality of the user-selected similarity measure, requires multiple hyper-parameters to manually tune, demands high computational resources and includes in the results only the classification performances together with the pathway names.

There is the need of new classifiers able to get the benefits of both the methodologies: classification and enrichment. As defined by Fabris et al., from the enrichment side the new method should be computationally light, easy to understand, provide more information than the probability value and not requiring assumptions to satisfy. While, from the classification side, it should be non-parametric, interpretable, consider multivariate interactions between features and patient classes, not requiring hyper-parameters difficult to set by the final user and cope well with both high-class imbalance and structured feature types.

We want to stand up to the challenge by proposing the pathway-based classifier called Simpati. Our method inherits the work done with netDx but evolves it to propose a totally new approach. It is the first method to use pathway-specific patient similarity networks and to adopt ad-hoc new strategies for processing this type of graph. It does not depend on hyper-parameters, measures, or external software that the user has to define or install. Simpati introduces novelties in every step of the classification based on PSNs, handles outliers, captures a unique paradigm of enriched pathways, uses a new recommender system to predict the class of an unknown patient and integrates graphical solutions to allow the visualization of the networks and the data generated during the workflow.

## 2. METHODS

### 2.1 OVERVIEW

Simpati considers the patient’s biological profiles (e.g., genes per patients) divided into classes based on a clinical information (e.g., cases versus controls). It prepares the profiles singularly applying guilty-by-association approach to determine how much each biological feature (e.g., gene) is associated and involved with the other ones and so to the overall profile. Higher is the guilty score and more the feature is involved in the patient’s biology. Simpati proceeds by building a pathway-specific patient similarity network (psPSN). It determines how much each pair of patients is similarly involved in the pathway. If the members of one class are more similar (i.e., stronger intra-similarities) than the opposite patients and the two classes are not similar (i.e., weak inter-similarities), then Simpati recognizes the psPSN as signature. If the classes are likely to contain outlier patients (i.e., patients not showing the same pathway activity as the rest of the class), then Simpati performs a filtering to keep only the most representative members of each class and re-test the psPSN for being signature. Unknown patients are classified in the best pathways based on their similarities with known patients and on how much they fit in the representative subgroups of the classes (i.e., more they are similar to the representatives of a class and more they fit). As results, Simpati provides the classes of the unknown patients, the tested statistically significant signature pathways divided into up and down involved (new pathway activity paradigm based on similarity of guilty scores), the patient similarity networks in vectorial and graphical format to allow further analysis of the subjects in studies using graph-theory methods, the guilty scores associated to the biological single-level features and all the data produced during the workflow.

### 2.2 DATA PREPARATION

Simpati works with patient’s biological profiles (e.g. gene expression profiles), the classes of the patients (e.g. cases and controls), a list of pathways and an interaction network (e.g. gene-gene interaction network). Simpati is designed to handle multiple biological omics but requires that the type of biological feature (e.g. gene) describing the patients is the same one that composes the pathways and the network which models how the features interact or are associated. In this study, we tested Simpati in the classification of Early versus Late cancer stage patients. In fact, identifying the cancer mechanisms which drive the tumor from early to late stages is challenging [31–33] but it can improve the early cancer diagnosis, lead to develop more precise therapeutic strategies and increase the survival rates [34]. A late-stage cancer spreads to nearby lymph nodes and other organs, the survival rate decreases due to the necessity of more advanced and risky treatment strategies. While, early localized stages are easier to treat and have better survival rates [33,35–37]. For setting up this biological and pathway-based classification challenge, we collected data about Liver hepatocellular carcinoma (LIHC), Stomach adenocarcinoma (STAD), Kidney renal clear cell carcinoma (KIRC), Bladder Urothelial Carcinoma (BLCA), Lung squamous cell carcinoma (LUSC) and Esophageal carcinoma (ESCA) cancers from The Cancer Genome Atlas (TCGA) using the R packages curatedTCGAData [38] and TCGAutils [39]. We kept only the patients having RNA sequencing (RNAseq) data, somatic mutation data and the histological type as clinical information. We added a new information based on the pathological stage attribute. We applied a binarization and labelled the stage I and II in Early, while the stage III and IV in Late based on the tumor/node/metastasis (TNM) system [40–43]. We proceeded with preparing the biological omics. We followed the workflow defined by Law et al. [44] for the RNAseq. Genes not expressed at a biologically meaningful level have been filtered out to increase the reliability of the mean-variance relationship. We removed the differences between samples due to the depth of sequencing and normalised the data using the trimmed mean of M-values (TMM) [45] method. While somatic mutation data have been converted into a binary data type, where a value equal to one indicates a mutated gene in a patient and zero otherwise. We ended up with two biological omics for five datasets with 14 LIHC (7 Early, 7 Late), 21 STAD (8 Early, 13 Late), 37 KIRC (24 Early, 13 Late), 45 BLCA (8 Early, 37 Late), 75 LUSC (60 Early, 13 Late) and 152 ESCA (91 Early, 61 Late) patients. The first four datasets to simulate wet-lab routine studies and the last two to have more precise classification performances [46]. We then collected the pathways and created the biological interaction network. We retrieved the biological processes from the major databases MSigDB [47], GO [48] and Kegg [49], while we used Biogrid [50] to model the biological feature’s interactions. A node represents a gene, and the edges are experimental and manually curated gene-gene interactions (GGi) (564,325 interactions and 26,433 genes).

Formally, given a set of features *FEA* = {*F*_1_, *F*_2_, *F*_3_,…, *F_A_*} with 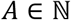 and of patient’s profiles *PAT* = [*P*_1_, *P*_2_, *P*_3_, *P_B_*} with 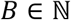 where each element is a vector of feature’s values, Simpati requires the concatenation of the patient’s profiles in a matrix *M*: *AxB* = (*m_a,b_*) where *m_a,b_* is equal to the value of *F_a_* in patient *P_b_*. The set of pathways *PATH* = {*PH*_1_, *PH*_2_, *PH*_3_…, *PH_C_*} where each element is defined by a finite set not exclusive of features (e.g., *PH*_1_ = {*F*_4_.*F*_5_, *F*_6_}). The biological network is defined as an undirected unweighted graph *GGi* = (*V, E*) where *V* = {*v*_1_, *v*_2_,…, *v_N_*} is a set of vertices (aka nodes) representing features *f*: *FEA* → *V* and *E* = {(*v_i_*, *v_j_*)|*v_i_*, *v_j_* ∈ *V*} is a set of pairs of vertices indicating edges (aka interactions, associations or relations) between features.

The adjacency matrix of *GGi* is the symmetric matrix *W*: *N* × *N* = (*w_i,j_*) defined by:

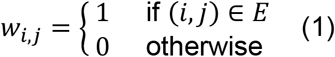

Finally, the patient’s classes are two vectors containing the indexes of the profiles belonging to them: 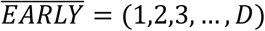 and 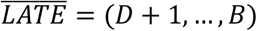 with *D* <= *B*.

### 2.3 NETWORK-BASED PROPAGATION

Simpati starts with protecting the privacy of the patients and enhancing the advantage of using the patient similarity network paradigm in the workflow. It converts the patient’s original information in anonymous labels. It then gives to the user the possibility to share data and results with or without the map. When the patient privacy is preserved, Simpati transforms the patient’s biological profiles using a network-based propagation algorithm. Each feature gets a new value based on its a priori information (e.g., expression or mutation value) and by its associations with all the other features. In other words, it gets a new value based on how much is found “guilty” of being involved in the overall patient’s biology and disease. This is based on three assumptions: the a priori information measures the strength of the link between the feature and the patient (e.g. the expression measures the gene importance and activity), the feature’s associations indicate shared molecular or phenotypic characteristics (e.g. interacting genes have similar cellular functions) [51] and a disease is rarely a consequence of an abnormality in a single feature but reflects the perturbations and signaling cascades provoked by multiple dysfunctional molecules [49].

In our application, Simpati maps the a priori values of the genes to their corresponding nodes in the GGi network. It propagates the values through the interactions using the propagation algorithm. Each node, even the one without value, gets a score which reflects its starting information and the amount given and received from its neighbours. The amount shared between nodes depends by the propagation type and the network topology. Simpati uses the random walk with restart (RWR) algorithm and the row-normalized version of the network. The RWR is a state-of-the-art networkbased propagation algorithm [52] and flexible standardization technique [51] which has been successfully applied for the disease characterization [53] and the prioritization of multiple disease-associated biological features as genes [52], pathways [54], miRNAs [55,56], lncRNA [57], proteins [58] and somatic mutations [59]. While, the row normalization guarantees that a node gives the same amount of information equally to all its neighbours independently by their degree [60,61]. This allows to not favour specific nodes against others.

Simpati uses the propagation to get the same continuous numeric datum from any input profile, to include in a single feature’s value also the perturbations provoked by the disease-specific activities of all the other patient’s molecules and to boost the signal-to-noise ratio [51]. For example, a poorly expressed gene gets a high score if close to strongly expressed genes and a non-mutated gene gets a high score if close to a mutated one. As consequences, this step allows to handle different biological omics (e.g. dense gene expression data, sparse somatic mutation data) [62–65], to use a novel ad-hoc similarity measure and to not let the interpretation and meaning of Simpati’s results depending by user-defined parameters.

For each profile *b* ∈ {1,…, *B*}, we define the set of its features represented as vertices with a priori information *SV_b_* = {*V_i_* ∈ *V* | ∀*i* ∈ {1,…, *N*} *s.t*. ∃ *F_i_* → *V_i_ and m_i,b_* ! = 0}. The RWR algorithm measures the importance of each node *v* to *SV_b_*. RWR mimics a walker that moves from a current node to a randomly selected adjacent node or goes back to source nodes with a back-probability *γ* ∈ (0, 1). RWR is described as follows:

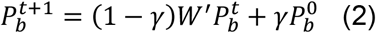

where *P^t^* is a *N* × 1 probability vector at time point *t* of which the *i_th_* element represents the probability of the walker being at node *v_i_* ∈ *V, P*^0^ is the *N* × 1 initial probability vector and defined as follows:

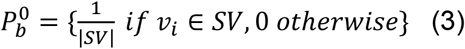

*W′* is the transition matrix of the graph GGi and 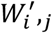 denotes a probability with which a walker at *v_i_* moves to *v_j_*. Formally, 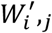 is defined based on the row normalization:

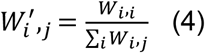

The propagated profile 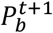 replaces the original profile 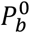 in the matrix *M*.

### 2.4 TRENDING MATCHING SIMILARITY MEASURE

Simpati works with patient’s propagated biological profiles. The feature has a score which measures how much is “guilty” of being associated and involved in the patient’s disease or generic clinical condition driving its biology with specific alterations and disfunctions. Higher the score and more the feature is involved. Plus, the propagation score is meaningful also between profiles. Lower the propagation score varies between patients and more the feature is assuming the same role. This point from a pathway perspective is important. Two patients may strongly involve one biological process but may act on it from different directions. For example, two patients may have high scores for the EGFR pathway genes (n=79) but have very different values for few ones as EGFR, JAK, IL-6 and GAB1. This may due to the fact that, the disease of one patient is acting on the pathway using exclusively EGFR and JAK [66], while the other is using IL-6 and GAB1 [67,68]. By literature, the direction of regulation of a pathway is disease specific and a biomarker.

We wanted to capture both the aspects of the propagation score in developing the similarity measure to estimate how much two patients were similar, so we designed a novel pairwise similarity measure called Trending Matching (TM) similarity (0 lowest, 1 highest). It is the weighted sum of two components: the mean and the variation of the propagation scores of the features belonging to a pathway. The first component measures how much the same feature is strongly or poorly involved in the patients. While the second component measures how much the same feature is similarly involved. For example, two patients described by a gene with high but different propagation scores (e.g., 1 and 0.7) are less similar than the pair which has lower but more close values (e.g., 0.8 and 0.7). The components are first determined for each single gene and then are summarized to represent the pathway. More the genes are strongly guilty, more the genes are similarly guilty and more the patients are considered similar in involving the pathway. This also prevents that, one outlier patient with genes strongly associated to the pathway can have a high similarity with another patient when they act on the process differently.

The trending matching similarity measures can be defined as follows; given a pathway *PH_u_* = {*F_a_* | *a* ∈{1,…, *A*}} and two patient’s profiles *P_b_* and *P_k_*, the similarity *TM_u_*(*P_b_, P_k_*) is the sum of three components:

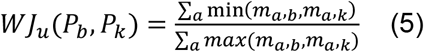

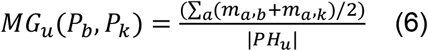

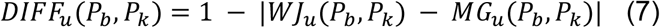

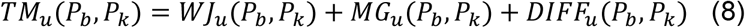

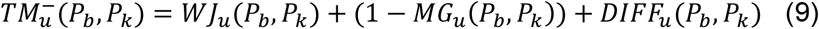

The TM similarity is designed in two variants. *TM_u_* is designed to capture what we will call upinvolved psPSNs, higher is the second component and higher is the similarity between two patients. The most cohesive class has higher propagation scores for the same genes than the opposite class. 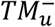 is designed to capture downinvolved psPSNs, lower is the second component and highest is the similarity between two patients. In the next sections and paragraphs, we will detail the operations using only *TM_u_* but the method performs every task with both the variants.

### 2.5 PATIENT SIMILARITY NETWORKS

Simpati aims to predict the class of an unknown patient comparing its propagated biological profile to the ones of the patients who are composing the classes of interest. For accomplishing the task, Simpati simulates a physician’s decision process applied to solve the diagnosis and prognosis of a new individual. Creation of a mental database of known patients linked by their similarity (e.g. Lung cancer patients and healthy controls), selection of the features in which patients of the same class are similar between each other but dissimilar from others (e.g. EGFR biomarker with overexpression in Lung cancer patients with respect healthy controls) and assessment of the clinical outcome of the new individual based on its similarities with the database ones.

Doing a parallelism, the mental database is a set of pathway-specific patient similarity networks. For each biological process, Simpati represents the patients as vertices of a network and weighs their interactions based on the Trending Matching similarity measure. Formally, the similarity network is defined as an undirected weighted graph *PSN_u_* = (*PV, PE*) composed by a set of vertices *PV* = {*pv*_1_, *pv*_2_,…, *pv_B_*} representing the patient’s profiles and a set of weighted edges *PE* = {(*pv_n_, pv_m_*)|*pv_n_, pv_m_* ∈ *PV*} with 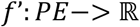 such that *f’*(*pv_n_, pv_m_*) = *TM*_*u*∈{1,…,*C*}_(*P_n_, P_m_*) representing how much the pair of patients is similar in a specific pathway *PH_u_*. The adjacency matrix corresponding to *PSN_u_*(*PV, PE*) is *W”*: *BxB* = (*w”_n,m_*) where *w”_n,m_* = *TM_u_*(*P_n_, P_m_*).

Simpati proceeds by selecting the pathways recognised as signature because represented by a PSN dividing the two classes while characterizing one. The members of one class must be more similar (i.e., stronger intra-similarities) than the opposite patients and the two classes not similar (i.e. weak inter-similarities). For this task, we developed a ranking system which evaluates a psPSN from 0 to 10 (the power of a PSN). Higher is the power and more a class is stronger than the opposite one and less the classes are similar (i.e., mixed together due to strong inter-similarities). First, we obtain three distributions based on the values of the similarities in the psPSN. The distribution of the intra-similarities possessed by the members of one class, the distribution of the intra-similarities of the opposite patients and the inter-similarities between the members of the two classes. For each distribution, we compute a low and high percentile (e.g., 0.4 and 0.6). Then, we check if the distribution of intra-similarities of one class has the low percentile greater than the high percentiles of the other two distributions. In case the condition is satisfied, we decrease the low percentile, increase the high percentile, and compare again. For example, power 7 is satisfied when the 20 percentile of the intra-similarities of one class is higher than the 80 percentile of the other distributions. The power 9 when the 15 percentile of one class is higher than the 85 percentile of the other distributions. When, a psPSN has at least power 1 is considered signature of the most cohesive patient class.

When the psPSN is built with the *TM_u_* similarity and has a power greater than 1 is considered signature and upinvolved because the members of the most cohesive class are similar due to higher feature’s guiltless than the patients of the opposite class. On the contrary, the psPSN built with the 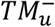 similarity is considered downinvolved because the most cohesive class has the lowest feature’s propagation values. Biologically speaking, the two classes are acting on the pathway differently, the members of one class are cohesive because their shared clinical condition is requiring and leading a precise alteration of the pathway, while the opposite class shows an heterogenous behaviour and a less need of acting on that cell function. We designed to capture this topological pattern and we do not require that the weak class must be cohesive following the study of Marquand et al. who reported that, assuming that both the classes in comparison are well defined precludes the inference of true diagnostic labels [69]. A clinical population may be composed of subjects belonging to different subtypes of the same disease or the heterogeneity may appear as result of misdiagnosis and comorbidities.

### 2.6 BEST FRIENDS CONNECTOR ALGORITHM

Simpati creates the database of psPSNs and then it aims to find the signatures. On the contrary of the starting situation in which a patient is described by the set of its biological feature’s values, now the patient is described by its similarities with both the members of its same class and the non-members. This is extraordinary with respect supervised enrichment (e.g., differential expression analysis and gene set enrichment analysis tools) and machine learning tools which do not normally neither expect nor assume that a patient can relate to the individuals of the opposite class. However, the patient similarity network paradigm intrinsically introduces the presence of outliers as it could be likely to have them in the study of a real clinical cohort [69] or disease-specific class [70,71]. The meaning behind is that the outlier patient shows a pathway activity which is different from the one exhibited by the rest of the members.

Simpati tackles this aspect assuming that if a psPSN is not signature, but it still includes a class more cohesive than the opposite one, then this psPSN is likely to contain outlier patients with low intra-similarities and high inter-similarities. If this case appears, Simpati performs a filtering to get rid of possible outliers which are misleading the topological analysis of the network.

We introduce the Best Friend Connector algorithm (BFC) for identifying the most representative and cohesive subgroup in each class, for removing members that are not similar to the majority and for maximizing the psPSN signature power (i.e., the grade of separability between the classes). At first, it determines which class is the strongest, then it performs the selection. For the strongest class, it finds the subgroup with the strongest intraclass, and weakest interclass similarities. For the weakest class, the subgroup with the weakest interclass similarities. The patients not selected in the subgroups are considered outliers and removed from the psPSN. Simpati keeps count of in how many pathways a patient has been considered outlier and it provides this information as result of the workflow to allow a further analysis of the a priori input data. The psPSN is tested for being signature and then recomposed as it was originally.

The algorithm exploits the concept of first order best friend (1BF), second order best friend (2BF) and outsiders. A patient is a 1BF of another member called root when their similarity is in the root’s best ones. A patient is a 2BF when it is 1BF of one root’s 1BF and it has the root as 1BF. An outsider is a patient that does not belong to the class which is undergoing the analysis. The algorithm performs the following operations for each class individually. It adjusts the weights of the intraclass connections. Precisely, it increases the similarity of two patients when both have a weak similarity with outsiders, while decreases in the opposite case. Iteratively, it considers one patient as root, it assesses the average of the intraclass similarities of the subgroup composed by his 1BFs and 2BFs. When each patient has been considered as root, the algorithm retrieves the set of best friends who got the highest average. This guarantees of avoiding selecting multiple strong subgroups identified by different roots which would not represent the starting class uniquely.

For explaining the pseudocode of the algorithm, we introduce the following mathematical details: Given four points in the Euclidian space *Q*(*x*1, *y*1), *E*(*x*2, *y*2), *R*(*x*3, *y*3), *T*(*x*4, *y*4), a vector of continuous values 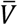, two indexes of patient’s profiles *b, k* ∈ {1,…, *B*} and a value *th* ∈ [1,…, 100], we define the formula of the quadrilateral area, the formula of the percentile, the average of the interclass similarities in two scenarios with respect *b* and *k*, and the first and second order best friend with respect *b*.

The quadrilateral area:

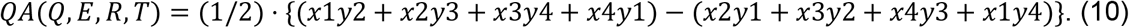

The percentile formula:

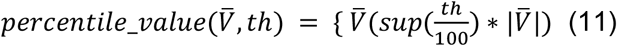

The average of the interclass similarities:

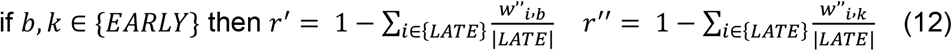

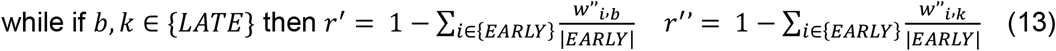

and the best friends with respect *b*:

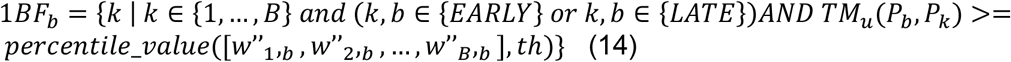

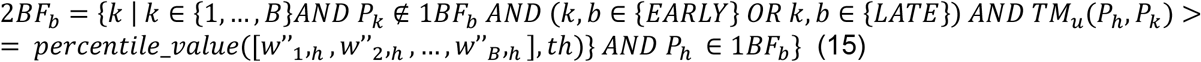

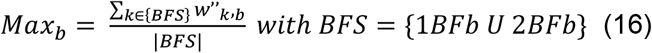

The Best Friends Connector algorithm is then shown in form of pseudocode in Fig.3 frame A.

**Fig.1.**
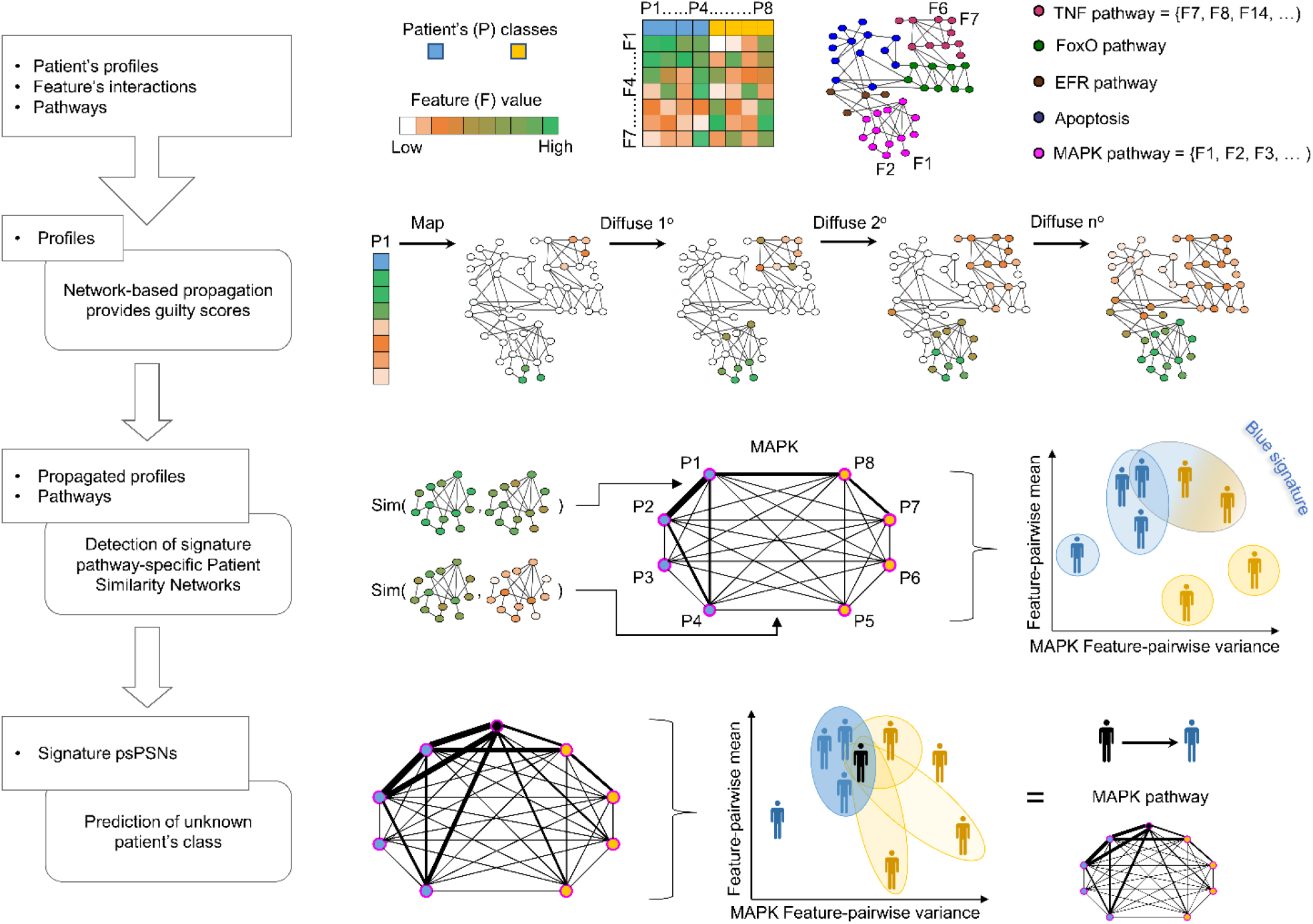
Workflow of Simpati. Patient profiles are divided in two classes and are described by biological features. A feature-feature interaction network together with pathways are further input data required by the software (e.g., gene-gene interaction network and KEGG pathways). All profiles are individually propagated over the network. The profile’s values are replaced by scores that reflect the feature’s starting information and interactions. Simpati proceeds by creating a patient similarity network for each pathway (psPSN). The pairwise similarity evaluates how much two patients have a similar pathway activity. It evaluates how much the features between two patients are close and high in term of propagation values. Two patients that act on a pathway with the same features and same expression values get the maximum similarity. In the figure, more an interaction is thick and more the two corresponding patients are similar. The psPSN is recognised as signature if one class is cohesive, one is sparse, and the two classes are not similar. In case of signature pathway, an unknown patient is classified based on how much is like the other patients.

**Fig.2.**
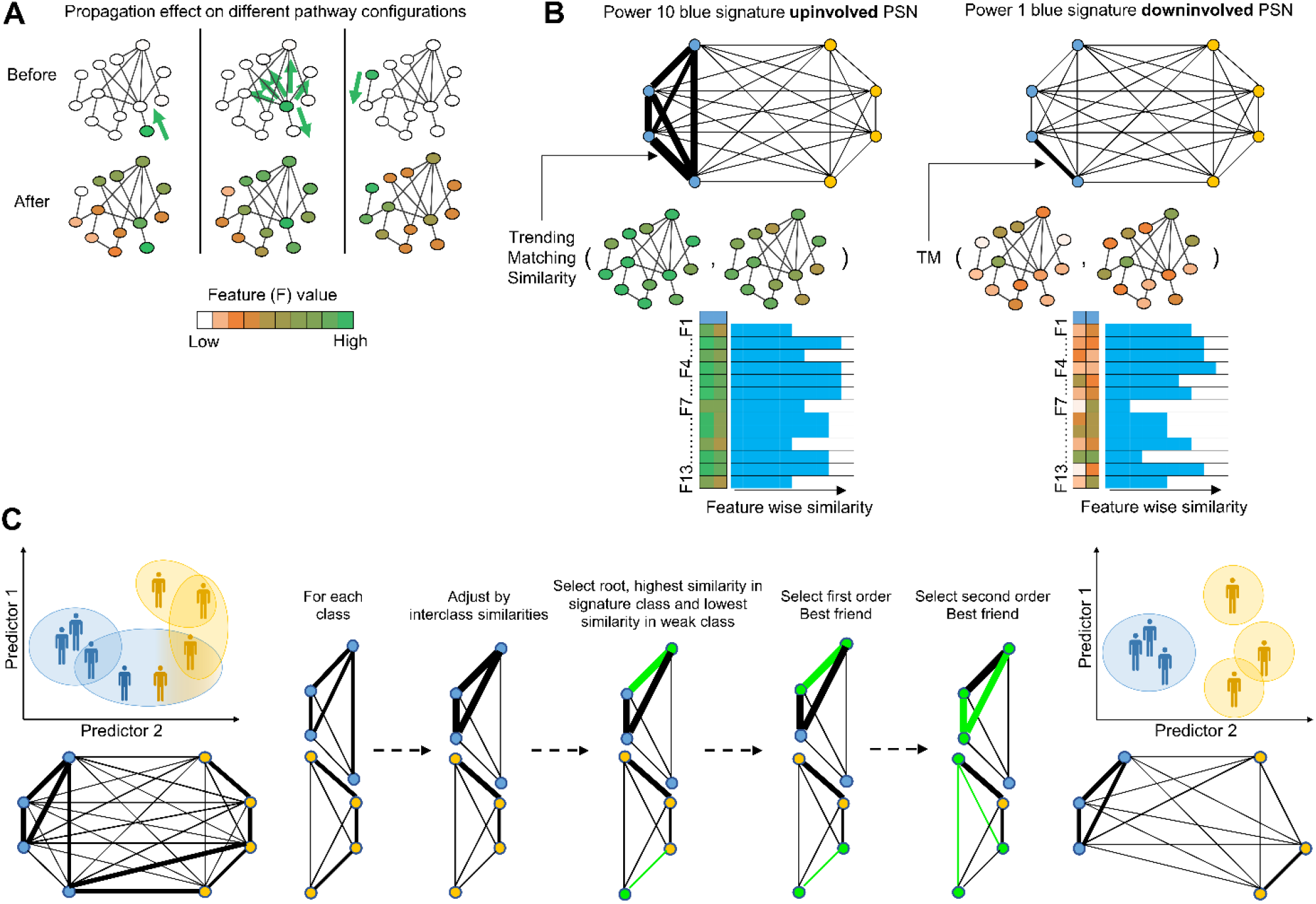
This figure includes graphical details about the phase of data preparation, creation of psPSNs and filtering of outliers based on the Best Friends Connector algorithm. The frame A shows how the result of the propagation changes due to the position of the patient’s molecule having a priori information. The propagation models how the pathway is specifically deregulated depending by which molecule is altered by the patient’s disease or generically active in a sample’s control profile. In the frame B, the Trending Matching similarity captures this aspect of the propagation. Two patients are considered strongly similar if each pathway’s feature is involved in the same way because it means that also the pathway is perturbated and deregulated similarly by the disease of the patients. Plus, the frame illustrates the difference between up and downinvolved pathway. In the first case, higher are the propagation scores and higher is the similarity between patients, while in the second case is the opposite. Frame C shows the steps which compose the Best Friends Connector algorithm. For each class, it adjusts the similarity between every pair of patients based on their connections with the members of the opposite class, then it selects first and second order best friends to compose the subgroup around a root. When all the patients have been considered as root once, the subgroup of best friends with the strongest interclass connectivity is chosen as best one.

**Fig.3.**
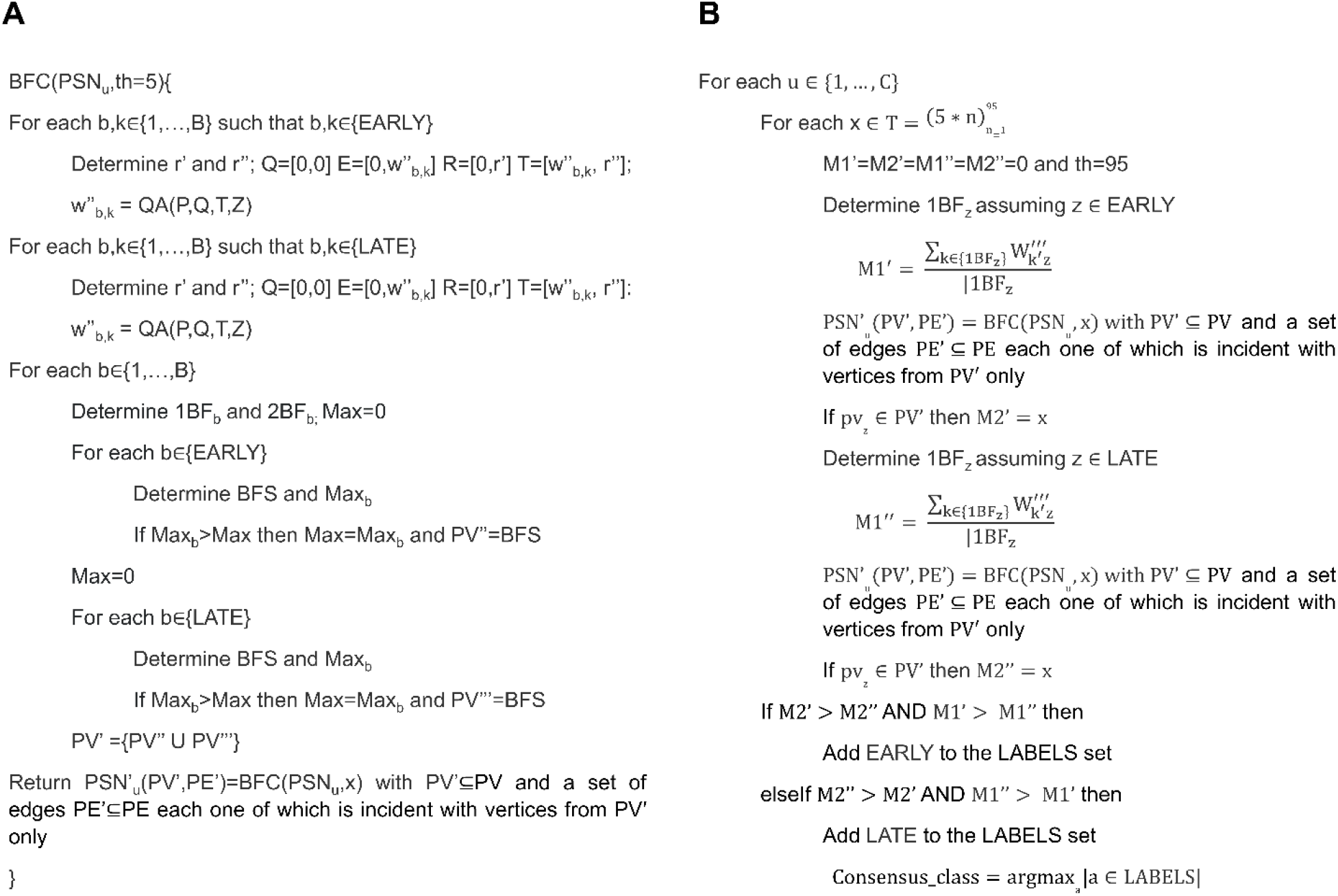
Figure of pseudocode relative to the two algorithms used in the Simpati implementation. The frame A shows the Best Friends Connector algorithm that is applied to a patient similarity network in form of adjacency matrix to filter outlier patients and detect the most cohesive subgroup. The frame B shows the step of class prediction related to an unknown patient performed at the end of the Simpati workflow.

**Fig.4.**
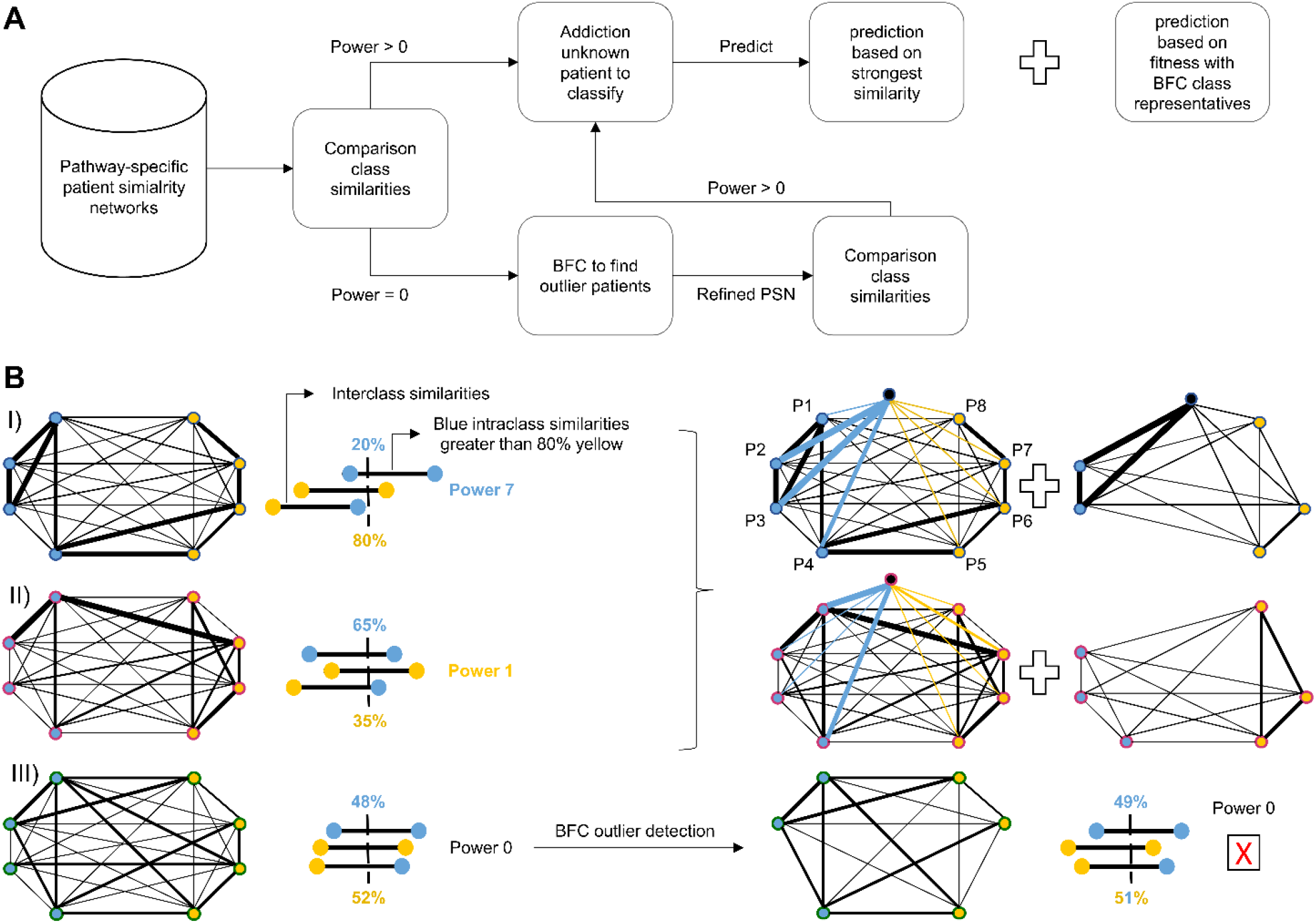
This figure shows the journey of a psPSN. In the frame A with a flow diagram, while in the frame B with different graphical scenarios. First, the psPSNs are created and compose a database. Then, Simpati determines the power of the networks. If a psPSN has a significative power is used for the classification, otherwise the method performs the BFC algorithm trying to detect outlier patients. If there is a cohesive subgroup in the psPSN and the removal of outliers significantly increase the power of the network, then the latter is reintroduced in the main workflow as signature pathway. The networks with sufficient power are used to classify an unknown patient. Simpati measures how much the patient is similar to the members of the classes and how much is distant from being an outlier. For this last step, Simpati runs the BFC algorithm iteratively getting each time a cohesive subgroup stronger than the one of previous iterations, more times the unknown patient is found in the best subgroup and less is considered outlier.

**Fig.5.**
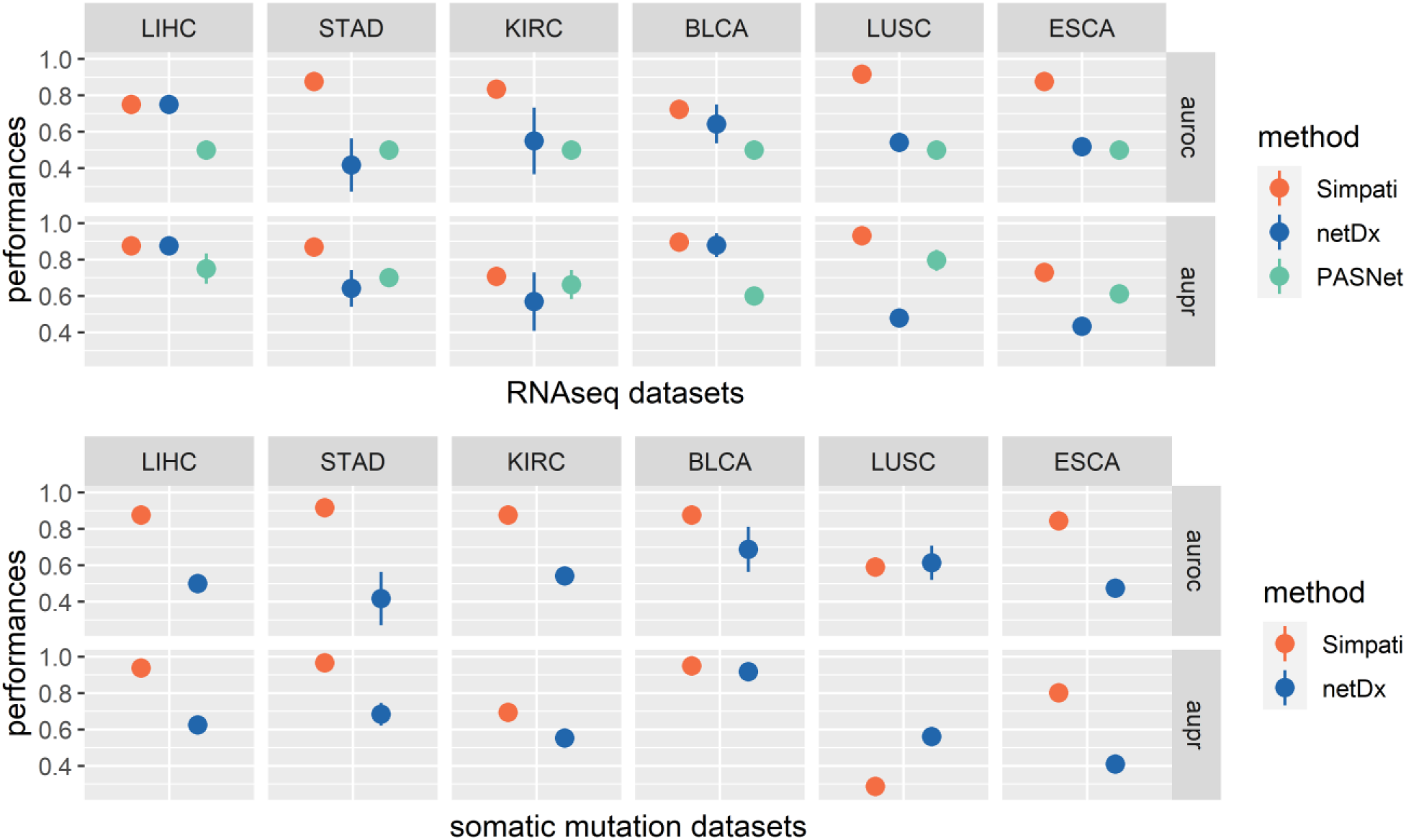
Comparison of the classification performances between the pathway-based classifiers Line plot of median (dot) classification performances with error bars (line). X-axis indicates the datasets. Y-axis indicates the value of area under the roc/pr curve. The same plot is presented twice, one including the performances when the methods classify the RNAseq data, while one the somatic mutations. PASNet does not have performances with somatic mutations because it does not handle sparse biological data. The plot shows that Simpati performs better than the competitors in all the datasets except for LUSC with somatic mutation.

**Fig.6.**
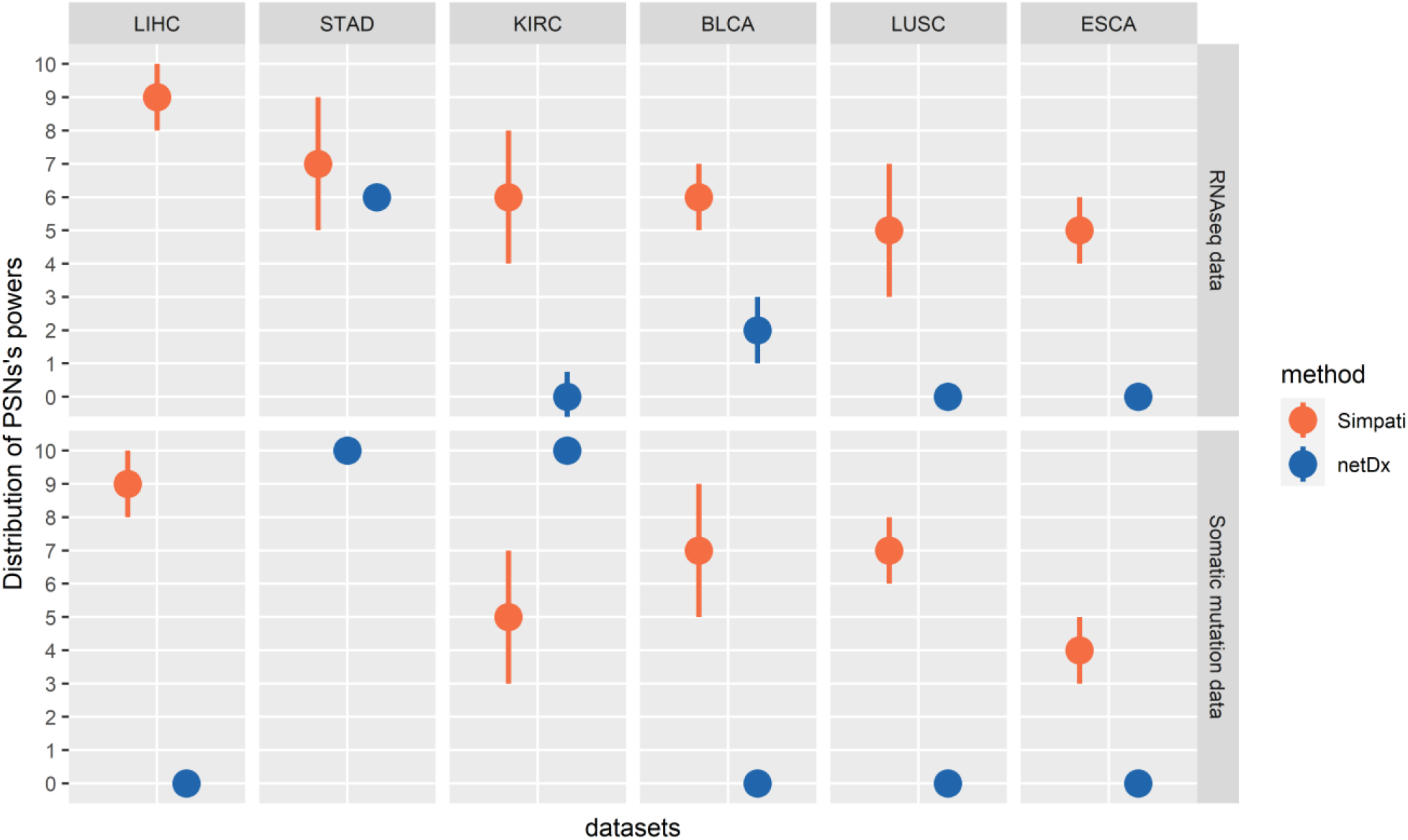
Comparison between the topology of the PSNs retrieved as result by netDx and Simpati with the TCGA datasets. The topology of the PSNs is measured with their power. Each frame of the image is dedicated to the pathways selected for the classification of a specific biological omic. The Y-axis indicates the power of the PSNs retrieved by a specific method. The X-axis indicates the datasets. Specifically, the dot indicates the median of the power of the PSNs resulted by applying a specific method with a specific dataset, while the line ranges based on the standard deviation of the same PSNs’ powers. Simpati selects better PSNs except in STAD patients described with somatic mutation profiles.

**Fig.7.**
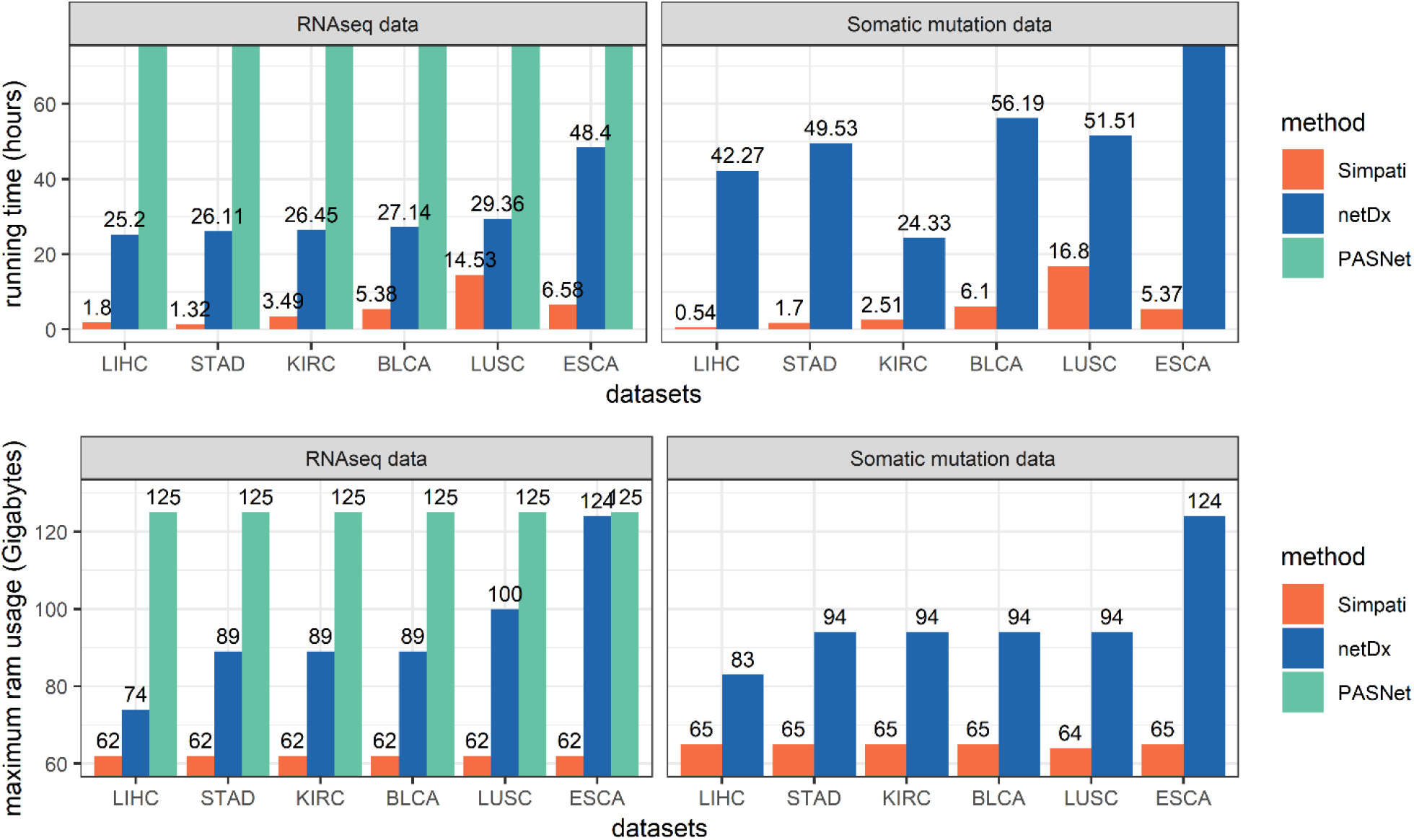
Barplot shows the comparison between the computational resources used by Simpati, netDx and PASNet to classify the TCGA datasets. The measures used for this comparison include the running time in hours and the memory ram in the maximum amount needed by the software in Gigabyte. In fact, the maximum amount is the real obstacle to the correct execution of the software. The X-axis indicates the datasets. The Y-axis indicates the measure. PASNet with the RNAseq data has a running time which exceeds the three days (72 hours). The plot shows how Simpati outperforms the competitors in time and memory for all the datasets.

**Fig.8.**
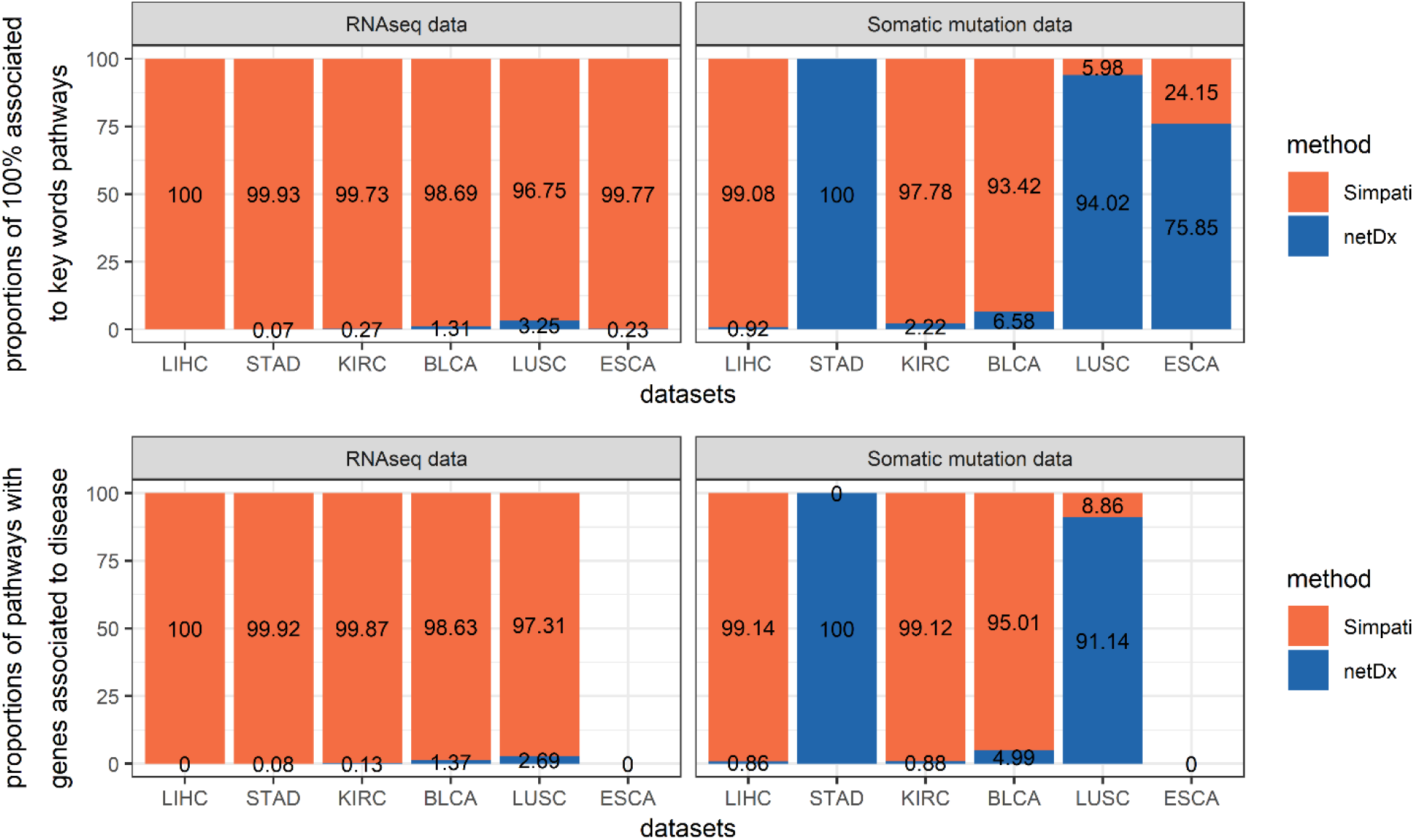
100% stacked bar chart shows the comparison between the enrichment statistics obtained by querying the resulting pathways of the different classifiers in Disgnet and the Human Protein Atlas with respect the patient’s cancer types. Each frame compares the methods based on how their pathways are qualified with a specific measure. The X-axis indicates the datasets in comparison. A bar is divided based on the number of pathways obtained by the two methods and satisfying the criteria indicated by the Y-axis. For example, netDx did not selected any pathway for the classification that satisfied the criteria in the classification of the LIHC patients with RNAseq profiles, while for the somatic mutation profiles Simpati has selected 99% more pathways.

**Fig.9.**
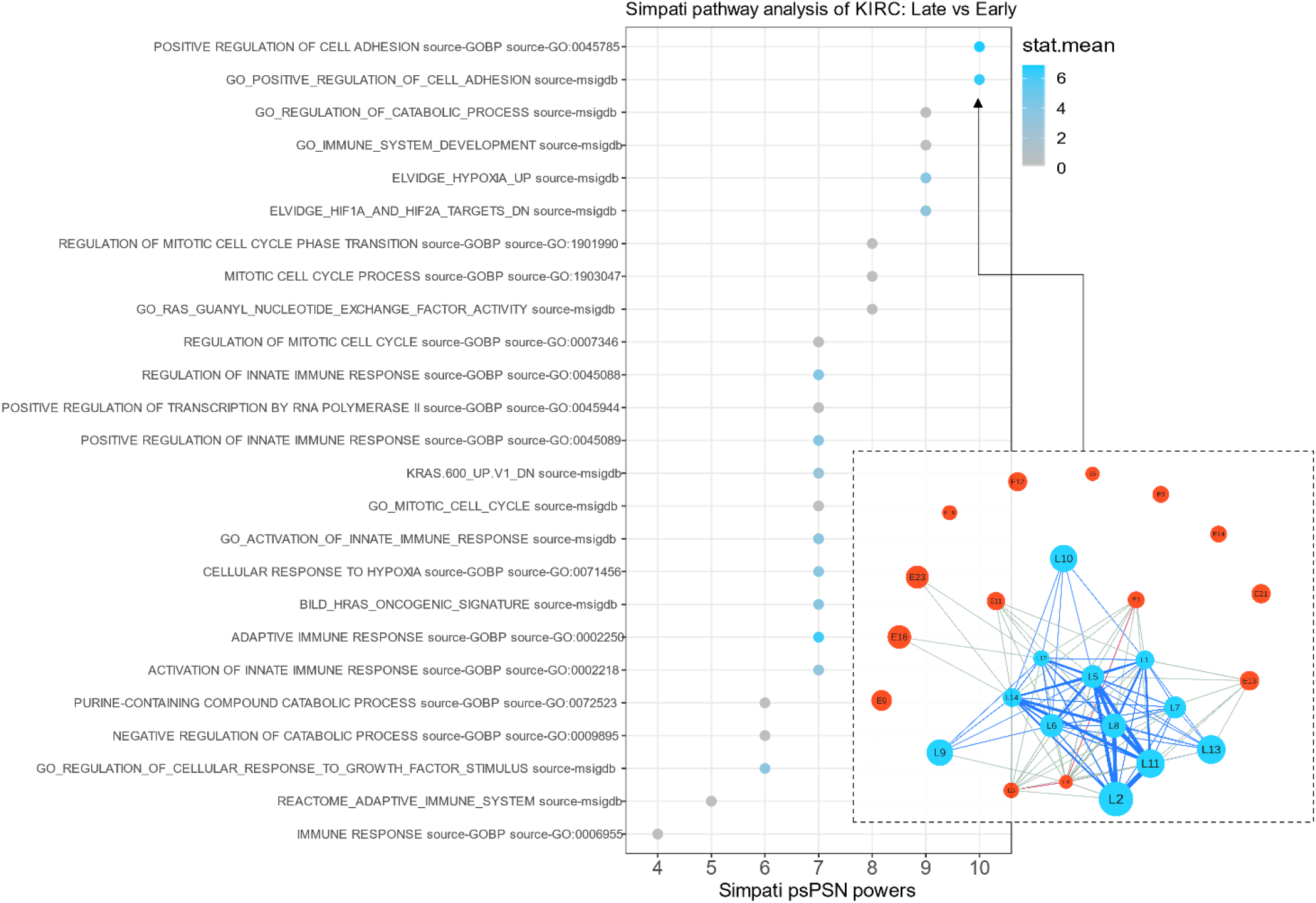
Dotplot showing the signature pathways detected by Simpati and the standard gene set enrichment analysis for the comparison of the KIRC patients between Late and Early stage. In the y-axis, there are the signature pathways and psPSNs resulting by performing Simpati. The x-axis indicates the power of the similarity networks. The legend describes how much the standard GSEA finds the same pathways altered between the two classes. The pathways with zero statistics mean (stat.mean) are not significant for the GSEA. Finally, in the bottom right cornet, the plot of the psPSN related to the positive regulation of the cell adhesion pathway. Blue nodes represent Late KIRC patients, while red nodes represent Early patients. The blue edges are the Late interclass similarities and the grey are the interclass.

### 2.7 CLASSIFICATION

Once Simpati created the database of signature psPSNs, it uses them to classify an unknown patient. This for understanding the quality of the selected pathways in characterizing and distinguish the classes in comparison. Simpati performs the operation continuing to follow the physician’s decision process. The unknown patient is compared to the ones already annotated in the mental database and assigned to the same class of who is most similar to. However, the only strength of similarity could be misleading. The unknown patient could have the strongest similarity with outlier members of the class. Therefore, we designed Simpati to consider also how much the unknown patient fits in the class.

The method prepares the unknown patient’s profile. The profile is replaced with its propagated version, compared to the patients in every signature pathway and added as new node in the corresponding psPSNs. Then, Simpati associates the profile to one of the classes based on the results of two approaches. For the first, it determines the average of the highest values of similarity that the patient has inside each class. The patient would be associated to the class with which has the highest average. While for the second approach, Simpati pretends that the patient belongs to one class and measures how much is far from being considered an outlier. The patient would be associated to the class in which is considered less outlier with respect the other members. In details about this step, the patient is simulated to belong to one class and the BFC algorithm is performed iteratively. At each run, the algorithm decreases the size of the subgroup of patients which retrieves. It stops when the patient does not belong to the best subgroup. Higher is the number of iterations and more the patient is considered having a stronger similarity with the class representatives than the other members which are more likely to be outliers. Simpati uses the iteration number as distance measure from the “outlier” status. Due to this, the patient would be candidate to be associated to the class in which survived the highest number of iterations.

Simpati associates the patient to the class that has been predicted by both the approaches. In case the results are not concordant, then Simpati does not make the prediction and the pathway together with its PSN are removed from the downstream operations. This step is performed for all signature psPSNs, then Simpati performs the consensus prediction. The patient’s definitive class is the one to which has been most frequently assigned.

Formally this would be, a new patient’s profiles *P_z_* such that *z* ∉ *EARLY and z* ∉ *LATE* is added as node in each patient similarity network. Let us define the new *PSN_u_*(*PV, PE*) composed by the set of nodes *PV* = {*pv*_1_, *pv*_2_,…, *pv_B_*, *pv_z_*} representing the patient’s profiles and the set of weighted edges *PE* = {(*pv_n_, pv_m_*)|*pv_n_, pv_m_*∈ *PV*} with 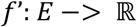 *s.t*. *f’*(*pv_n_, pv_m_*) = *TMu*(*P_n_, P_m_*). The class of the new profile is found by a topological analysis of all the psPSNs shown in form of pseudocode in the figure 3 frame B.

### 2.8 OUTPUT

Simpati provides the signature pathways used in the classification step, the corresponding PSNs in vectorial format and reports statistics to allow further analysis and considerations. For example, the average of the intra and inter similarities for recognising the most cohesive class, the psPSN power to get the pathways which more separate the classes in comparison and a probability value (p.value). The latter is assessed testing the psPSN to retrieve an equal or higher power than the original one when patients are permutated between classes. This information allows to filter out pathways which have been detected as signature due to random.

Simpati also includes two tools for the visualization of the data produced with the workflow, one tool is an internal function able to produce a compact representation of a psPSN, while one is a graphical user interface (GUI) for the exploration of a patient’s propagated biological profile. The function provides a compact representation of a psPSN by reducing the patients which are visible as nodes. This is necessary to allow the user to understand how patients are similar between each other. In fact, more the number of nodes increases and more becomes difficult to follow the edges of the network and so how the patients are similar between each other. To do so, Simpati groups up patients of the same class that are considered similar, chooses a patient to represent each subgroup and filters the original network by keeping only the chosen (i.e., the representative of each subgroup). First, it determines how much every pair of patients are similar in the network. It uses the measure *WJ_u_*(*P_b_, P_k_*) which is applied between two patient’s profiles composed by their similarity values in the psPSN. It gets a psPSN of the psPSN (aka psPSN^2^). Simpati proceeds and iteratively performs the BFC algorithm on the psPSN^2^. In this case, the BFC algorithm is not applied to filter outliers but to detect multiple cohesive subgroups. At every iteration, the BFC algorithm detects and removes the most cohesive subgroup composed by the twenty percent of the original members of one class. When the BFC cannot be applied anymore due to the small size of the class, Simpati filters the original psPSN and keeps only the class members which have not been grouped in the last iteration together with the root of each class-specific detected cohesive subgroup. Further about the aesthetic aspects, the size of a node is used to indicate how much the relative patient is similar inside its own class compared to how much is similar with the outsiders. This is assessed with the difference between two PageRank [72] centrality scores, one is measured only with the patient connected by similarity to the members of its class and one with only the outsiders. Higher the difference, higher the size of the node, more central the patient is in its class and less similar to the outsiders. While, the position of the nodes in the plot of the psPSN is determined using the Fruchterman & Reingold’s force-directed layout [73]. The network becomes compact, easy to analyse and still representative of how all patients are similar. Thanks to the plot, the user can understand if the psPSN is correctly a signature, see how the similarities are distributed, identify which patient is crucial for the connectivity of its own class and which is instead behaving as outlier.

Complementary to the visualization of the psPSN, we also provide an R shiny graphical user interface (GUI) to allow the exploration of the propagation effect over a patient’s profile. This enables the user to understand how the values of the patient’s biological features changed and for which reason. For example, a gene with low expression value that has been removed from the Limma analysis due to the function “filterByExpr” in the differential expression analysis, it may get a high propagation score and the user may get interested in understanding the reason. We believe that this can be another useful instrument to make the method and the data more accessible.

### 2.9 ENRICHMENT

The power, the p.value, the distribution of the similarities are all technical information regarding a psPSN that allow to understand how patients and classes are structured. However, they offer a limited utility in prioritizing the best pathways because they are not related to any biological background of the patients. For this reason, in case the patient’s features are genes, we designed Simpati to perform a query in the Disgnet [74] and Human Protein Atlas [75] (HPA). DisGeNET is a database which provides open access to annotated disease associations with genes and variants. While HPA is a unique world-leading effort to map all the human proteins in cells, tissues, and organs in the human body using antibody-based imaging, mass spectrometry-based proteomics, transcriptomics, and systems biology. Simpati requires the semantic type of the patient’s disease (e.g., Neoplastic Process, Congenital Abnormality, etc..) and key words (e.g., TCGA-KIRC: Kidney, Renal, Carcinoma). Then, it gets which published articles have associated the genes of a signature pathway to the semantic type and to the key words. Plus, in case of cancer patients, this integration allows to get which genes are favourable to be prognostic, while in case of non-cancer disease which genes are associated to the user-defined tissue. As indicated by Lin et al. [76] this operation allows to prioritize the signature pathways based on their associations with the patient’s clinical outcome and to understand better the validity of Simpati results.

### 2.10 WORKFLOW OF TESTING

Simpati ability to classify the classes in comparison is tested with a leave one out cross validation (LOO-CV). Given a dataset of patient’s biological profiles and the classes associated to them, Simpati iteratively performs the following operations: one patient is considered unknown and compose the testing set, while the remaining patients are considered known and used as training set. The latter is used to build the psPSNs, to find the signature pathways and as ground truth in the classification step. While, the testing patient has the biological profile which class must be predicted. In the end, the predicted classes of the testing patients collected from all the iterations are compared to their real ones for determining the classification performances. Simpati is designed to value its ability to predict based on two measures following netDx design [30]. The first one called AUC-ROC is the area under the curve where the x-axis is the false positive rate (FPR) and the y-axis is true positive rate (TPR). While the second one called AUC-PR is the area under the curve where the x-axis is the recall and y-axis is the precision [77].

## 3. RESULTS

### 3.1 CLASSIFICATION COMPARISON

We tested Simpati performances to classify patients of five different TCGA cancer types described by two biological omics: transcriptomics with gene expression data and genomics with somatic mutations. The classes assigned to the patients were Early or Late based on their cancer stage. We increased the challenge including the performances of the current published generic-purpose pathway-based classifiers: netDx [30] and PASNet [21]. netDx creates a database of predictive PSNs associated to pathways for each class, builds a consensus similarity network and applies GeneMANIA (state-of-art gene function prediction algorithm) for the prediction of the testing patients. netDx tests its performances with a 10-fold cross-validation which in each run includes another cross-validation for the selection of the most predictive PSN. While, PASNet incorporates biological pathways in a Deep Neural Network. The neural network is composed by a gene layer (an input layer), a pathway layer, a hidden layer that represents hierarchical relationships among biological pathways and an output layer that corresponds to the patient classes. PASNet tests its performances with a stratified 5-fold cross-validation repeated 10 times. The two competitors either support or use the classification evaluation based on the area under the receiver operating characteristic curve and the area under precision-recall curve measures and they differ from canonical supervised machine learning algorithms. For these reasons, we performed the comparison using each method based on how it has been designed and following the vignettes provided by the authors.

Simpati performs better than the competitors for both the measures with all the omics. Simpati also proves to be more reliable in each dataset with a standard error equal to zero due to its leave one out cross-validation approach. While the performances of the competitors highlight common classification issues; their performances vary a lot probably due to the number of patients, the size of the classes in comparison and the ability of the classifier to naturally handle multiple omics and data types.

### 3.2 SIMILARITY NETWORK COMPARISON

Simpati and netDx provide pathways and related PSNs as result of the analysis. However, the methods use different techniques for the pathway selection. netDx selects a pathway if the corresponding PSN allows GeneMANIA to correctly predict the classes of the training and testing patients. While Simpati selects a pathway if the corresponding PSN topologically separates the classes in comparison. The resulting pathways and related PSNs should help to characterize the patient classes, explain why they have been used to predict and they should increase the interpretability of the model. We compared the topology of the PSNs selected by the two contenders based on their power.

Simpati provides more pathways with high power than netDx in all the datasets except one. This is probably due to how the selection is done. Simpati discerns PSNs based on their topology and then performs the classification. While netDx evaluates a pathway based on the mere ability of the GeneMANIA algorithm to use its PSN for classifying. This makes the difference in terms of interpretability of the model. From the final user perspective, Simpati’s psPSNs together with their visual representation make easier to understand why they have been selected for the classification and can be perceived as more trustable.

### 3.3 COMPUTATION RESOURCES COMPARISON

The patient similarity network paradigm used by Simpati and netDx brings many advantages both in the feature selection, in the classification phase and in the overall interpretability of the software. However, these pros come with a price which is the software scalability already introduced as challenge by Pai et al. [22]. A PSN is a complete graph that the methods build with all the patients and for every pathway. This means that an increment in the number of patients and in the number of annotated pathways lead the methods to require more computational resources. netDx and Simpati faced this point with different approaches. netDx is implemented in R and Java, uses the disk to save temporary files and applies a sparsification of the PSNs to decrease the number of edges and so the amount of information associated to them. While Simpati is implemented completely in R, natively support parallel computing and handles all the data of the workflow as sparse matrices or vectors. To understand which software handles better this issue, we captured the ram usage and the running time that each method required to classify the TCGA datasets with the same hardware setting (AMD Ryzen Threadripper 3970X 32-Core Processor, 251 Gigabyte System memory and Linux ubuntu-1804-slurm 5.4.0-72-generic). Simpati resulted to be more efficient in both the running time and the ram used.

### 3.4 ENRICHMENT COMPARISON

Pathway-based classifiers aim to classify correctly unknown patients using the biological pathway information. This means that the prediction of a patient’s class passes through the selection of pathways which due to method-specific criteria are considered useful for the task. In a cross-validation setting, the final classification performances indicate how much the classifier is reliable and better than a random predictor. However, they do not represent a measure of how much the pathways are biologically significant. A classifier as Simpati can provide further details about how it used the pathways, why it selected them and the biological interpretation under the filtering criteria but still, this information does not allow to understand if pathways are biologically meaningful. For this reason, we designed Simpati to integrate an enrichment step and we performed this operation also to the results of the other competitors. Precisely, we kept only the resulted significant pathways having at least one publication associated to each of the key words defined per dataset and having at least the 90% of the genes associated to the patient’s specific cancer. Then, we compared the numbers of pathways satisfying these constraints.

This analysis highlights that Simpati is both able to select, use and provide biologically significant pathways directly associated to the patients that it is classifying and that performs better than the competitors. netDx retrieves always much fewer pathways biologically associated to the tumor of the patient’s profiles than Simpati. We have been unable to include PASNet due to its lack in providing the pathways that it considers significant and predictive of the patients classified.

### 3.5 PATHWAY ANALYSIS

In this last section, we compare the Simpati signature pathways obtained from the classification of KIRC patients with literature findings. We considered the key pathways identified by Pang et al. [11] and Cui et al. [10].

Late stage KIRC regulates the cell adhesion to escape from the immune system. It disturbs the ATP supplement, making cells to adopt anaerobic respiration, producing an ATP deficit and hypoxia. It provokes an immune response from the host organism, regulates the PI3K-Akt signalling pathway, promotes anabolism and inhibits catabolism.

Simpati has been able to find key pathways differentiating the pathological stages of the patients. Specifically, it confirmed the relationship between the late cancer stage and the upregulation of the cell adhesion, response of the immune system and hypoxia. For this comparison, we also included the results of the standard gene set enrichment analysis (GSEA) performed thanks to the GAGE method [78]. The two methods found similar key pathways. However, GSEA provided the statistics mean, while Simpati provided the patient similarity network.

## 4. CONCLUSIONS

We propose the pathway-based classifier called Simpati. The method can be applied to different omics, proved to obtain quality classification performances and detect signature pathways. In other words, it identifies biological processes that distinguish and uniquely characterize the clinical classes of the patients in comparison.

On top of the technical conclusions, we want to suggest Simpati as tool for computational biologists and bioinformaticians that want to get unique insights about their patients or samples. We designed Simpati to simulate a physician’s decision process applied to solve the diagnosis and prognosis of a new individual. As a physician, our software processes, stores and learns information related to the patients. All the data used during the classification are then made available for allowing further analysis.

Simpati associates to each single biological feature (e.g. gene, protein, mutation, …) a propagation score which reflects the overall biology of the patient. A high score indicates that the feature is strongly involved in the patient’s biology, while a low score the opposite. The scores can be explored in two ways. They can be considered as values of a standard high-dimensional matrix with patients at the columns and features at the row. They can be visually taken into account in an ad-hoc graphical-user interface. The matrix format allows any statistical analysis with clusterProfiler [79], while the GUI permits to understand how much specific biological features of interest are important without any programming and statistical knowledge. The information retrieved by analysing the propagation scores can be combined to the results obtained from a differential expression (DE) analysis. For example, a DE list can be filtered to keep only the genes that have a high score in order to reduce the false positive or can be expanded integrating those genes that are DE in term of propagation values.

Simpati models pathways as patient similarity networks. In a psPSN, patients are connected to others based on how much their biology similarly regulate a specific pathway. Like in a social network in which people are connected to others based on their hobbies and how they practice them (e.g., the place, the effort, the time). More two patients involve and regulate similarly a pathway (e.g., with the same genes and with the same expression values) and more they are strongly connected. In case a pathway is found significant and of interest, it can be explored in two ways. The adjacency matrix or the graphical representation of the related psPSN. The matrix format allows any network analysis with NetworkToolbox [80], while the plot permits to have intuitions about how much the patient classes separate and to identify patients that are central or tend to be outlier. The information retrieved by analysing the topology of the psPSNs can be used to verify the clinical information associated to the patients, identify subclasses, and can be combined to the results obtained from a clustering analysis or a non-negative matrix factorization (NMF) [81]. For example, the subclasses of patients that have been identified with an unsupervised technique can be checked in the psPSNs to find in which pathways are mostly similar.

Simpati finds signature pathways to characterize and distinguish the patient classes in comparison. The pathways must satisfy a constraint. The members of one class must be more similar than the opposite patients. Then, it uses the similarities to predict the class of new patients. In this sense, Simpati can be combined to a standard gene set enrichment analysis because detects pathways that satisfy a criterium not taken into account by other tools.

Simpati does not assume that the patient classes are well defined, and it considers the possibility that members of the same class may regulate the same biological process differently. When this is likely to happen with a pathway, Simpati identifies the patients that most represent their own class, uses only them to check the signature condition and the remaining members are considered outliers. For this reason, Simpati is suitable to real case scenarios which often include either patients or samples associated to clinical outcomes due to a priori information by wet lab scientists. The latter check the expected sample classes using a principal component analysis or clustering. However, both the methods are not designed to detect differences at the level of pathways or single biological features which could reveal unique biological aspects of a sample and differentiate it from the rest of its class. For example, in a knock-out study, samples are labelled as knocked-out based on the experiment but this does not necessarily imply that each member of the clinical population shows changes in gene expression levels against the control group. Standard gene set enrichment analysis tools are ineffective due the possible low variation between the classes and possible knock-out samples not showing any change.

## 5. DISCUSSION

Generic purpose pathway-based classifiers propose themselves as powerful tools for classifying patients and providing biologically meaningful results in form of pathways. The first one has been introduced in the 2010 but, at the time of writing, there remain very few. This is due to the many challenges that must be faced to produce classifiers able to combine biological omics and pathways, while being able to compete with standard enrichment tools. We tried to report and detail all the issues related to the development of this kind of machine learning algorithm. At the same time, we tried to build a software that could have been considered a future example for other researchers. Thanks to the combination of new and popular strategies, Simpati proves that is possible to both obtain satisfying results and tackle common issues in the pathway-based classification.

The preparation of the patient’s biological profiles with a transformation technique as the network propagation allows to get the same kind of data and information before the classification. This allows the researchers to develop a workflow which is flexible, consistent, and involving less hyper-parameters. As a matter of fact, we developed the Trending Matching similarity to capture a specific relationship between the patient’s propagated profiles and the scored genes. On the contrary, netDx suggests using the Pearson correlation as default measure directly on the raw profiles but the authors did not provide a biological interpretation of what kind of patient similarity leads to catch and leave the user to the uncontrolled intrinsic disadvantages of the measure [82]. For example, the correlation is biased and leads to incorrect inferences when considers genes that have not been perturbed (e.g., environmentally or genetically) in order to cause a meaningful change in expression level [83].

The selection of the predictive pathways including criteria that can be explained from a biological point of view allows the classifier to not drift away from the patient’s biology. For example, both PASNet [21] and netDx [29] select the pathways which perform the best in predicting the training patients. This approach is undeniable well suited for securing the ability to predict an unknown patient. However, it may lead to select pathways which are useful for the algorithm of prediction and meaningless for the patient’s biology. On the contrary, Simpati performs a first selection based on criteria that can be biologically interpretated and then it analyses the pathways for the classification. As we proved with our results, a biological selection does not necessarily negatively affect the final classification performances but it indeed changes positively the resulting pathways produced by the classifier.

The analysis of outlier training patients because their biological features are not showing the same activity in a pathway as the rest of the class increases the granularity of the cell process description that the classifier handles and provides as result. This brings multiple advantages. It makes the classifier less sensitive to how much the data have been cleaned, how the patient classes have been defined, and it allows to give a hint about subclasses of patients which are using different pathways at the net of one shared clinical status.

At the same time, it is worth to also mention the price of such strategies. The propagation leads the classifier to require also the network of interactions or associations between the features of the biological omic. The selection of pathways based on criteria which are biologically explainable is not trivial and may makes the classification inconclusive due to no pathway passing the filter. The analysis of the outliers requires either parameters to set up which may be not correct for all the applications or hyper-parameters to determine.

## 6. AVAILABILITY OF DATA AND MATERIALS

All the work has been made in R programming language, from the data extraction to the enrichment. We provide a github repository with a tutorial about how to replicate all the results of this project: https://github.com/LucaGiudice/supplementary-Simpati. We provide an R package to use Simpati: https://github.com/LucaGiudice/Simpati and to use the GUI: https://github.com/LucaGiudice/propaGUIation

## AUTHOR INFORMATION

LG designed and developed the software. LG wrote the article and supervised the project. LG produced the images and created the github repositories.

## ACKNOWLEDGEMENTS

I want to thank my best friend Samuele Cancellieri who supported me during the development of this work, who has been the true reader and critic of this manuscript, and that was always there to hear my difficulties and complains. I want to thank Shraddha Pai, author of netDx, who inspired me to develop this software and taught me what to improve and how to do always better. I want to thank Gary Bader who started this adventure of mine by letting me work on patient similarity networks in his research group during my master thesis. I want to thank my girlfriend Lucy who had to deal with all the stress of mine caused by this project.

